# Super-Resolution Processing Enables *In Situ* Subtomogram Averaging Beyond the Nyquist Limit

**DOI:** 10.64898/2026.10.17.759810

**Authors:** Euan Pyle, Sultan Bekbayev, Vinolia Dmello, Simone Mattei

**Affiliations:** Molecular Systems Biology Unit, European Molecular Biology Laboratory, Heidelberg, Germany; Cell Biology and Biophysics Unit, European Molecular Biology Laboratory, Heidelberg, Germany; B CUBE - Center for Molecular Bioengineering, TUD Dresden University of Technology, Dresden, Germany; Faculty of Biology, TUD Dresden University of Technology, Dresden, Germany; Cluster of Excellence Physics of Life, TUD Dresden University of Technology, Dresden, Germany; EMBL Imaging Centre, European Molecular Biology Laboratory, Heidelberg, Germany

**Author notes:** Correspondence should be addressed to Simone Mattei.

## Abstract

*In situ* subtomogram averaging (STA) for high-resolution structure determination requires fine pixel sampling, forcing a trade-off between achievable resolution and the field of view (FOV) recorded in each tomogram. Electron-event representation (EER) data can be rendered at half the physical pixel size, but whether super-resolution processing can recover useful information beyond the physical Nyquist limit from noisy *in situ* data remains unclear. Here, we show that super-resolution processing of EER data recovers biologically interpretable information beyond a physical Nyquist limit of 4.78 Å in two *in situ* datasets. STA of unidentified virus particles in a *Drosophila melanogaster* larva yielded a 3.42 Å inner capsid reconstruction, enabling *de novo* model building and identification of the particles as Drosophila A virus. *In situ* STA of ribosomes in *Saccharomyces cerevisiae* reached 3.18 Å, with local resolution extending to 2.78 Å. EER acquisition therefore provides greater flexibility in choosing the acquisition magnification: existing datasets can be re-rendered at the super-resolution pixel size when physical sampling becomes limiting, while lower-magnification acquisition can be considered when a larger FOV improves target capture or cellular context.

## Introduction

Cryo-electron tomography (cryo-ET) combined with subtomogram averaging (STA) can determine high-resolution structures of macromolecular complexes directly in cells and tissues, preserving their near-native molecular environment ^1–13^. However, sub-nanometre reconstructions commonly require the alignment and averaging of thousands of particles from datasets containing large numbers of tilt series ^1,2,14^. For cellular cryo-ET projects, the achievable dataset size is often constrained by the time required to prepare electron-transparent lamellae and acquire tilt series ^15,16^. Cellular lamellae are challenging to produce ^17,18^, especially for high-pressure frozen specimens which require complex lift-out or waffle workflows ^19–22^. Therefore, maximising the number of useful particles in each collected tilt series is important to minimise the investment of human and hardware resources dedicated to the structural determination of the macromolecular complex of interest.

The magnification to be used for data acquisition is therefore a consequential experimental choice. Imaging at lower magnification increases the field of view (FOV), providing more cellular context and, in many cases, more particles per tomogram. A larger FOV can also cover a given lamella area with fewer tilt series, minimising the microscopy time required for imaging the full lamella. However, the selection of a lower magnification comes at the cost of a larger physical pixel size, the size of one detector pixel at the specimen. As the Nyquist limit is twice the pixel size, lower-magnification acquisition restricts the resolution attainable by STA. Researchers must therefore balance particle yield and contextual information against the risk that the chosen sampling will limit the final reconstruction.

Direct-electron detectors (DEDs) provide a way to relax this trade-off. Each electron striking the detector deposits charge across several neighbouring pixels, allowing the position of the electron-event to be estimated with subpixel precision ^23–25^. DEDs commonly used in modern cryo-ET, such as the Gatan K2 and K3 as well as the Thermo Fisher Scientific Falcon 4 and 4i, conventionally process this signal in ‘counting’ mode, in which the electron-event is assigned to the pixel receiving the largest share of the charge distribution (Fig. S1). DEDs can also leverage the subpixel localisation precision of the electron-events to produce images at ‘super-resolution’ pixel size, typically half the physical pixel size. Gatan K2 and K3 detectors record super-resolution data in a dedicated acquisition mode ^24^, whereas Falcon 4 and 4i detectors store individual electron events in electron-event representation (EER) files on a 4 × 4 subpixel grid (Fig. S1) ^25^. EER data are rendered on this grid and Fourier-cropped to the chosen pixel size, so images can be processed at either the physical or a super-resolution pixel size. Rendering at the super-resolution pixel size doubles the sampling frequency. A target sampling can therefore be reached at half the magnification otherwise required, doubling the linear FOV and quadrupling the imaged area.

However, finer sampling does not necessarily translate into recoverable structural information. The detective quantum efficiency (DQE) decreases above the physical Nyquist limit, where each electron contributes less signal relative to noise ^26,27^. Super-resolution processing has nevertheless enabled reconstructions beyond the physical Nyquist limit in single-particle analysis (SPA) of purified specimens ^25,28,29^, where the larger FOV yields enough particles to offset the reduced DQE. Lamellae are thicker and more crowded than SPA specimens, so whether enough signal survives at these frequencies is uncertain. A previous STA attempt on purified apoferritin found no benefit ^30^, and recovery of information beyond the physical Nyquist limit has not been shown *in situ*.

Here, we show that processing Falcon 4i EER data at a super-resolution pixel size supports *in situ* STA beyond the physical Nyquist limit in two distinct datasets. Virus particles in lamellae prepared by serial lift-out from *Drosophila melanogaster* tissue yielded a 3.42 Å reconstruction from 20 tomograms acquired with a physical Nyquist limit of 4.78 Å, resolving side chains and enabling automated model building and the identification of the particles as Drosophila A virus. In a second, asymmetric test case, *Saccharomyces cerevisiae* ribosomes reached 3.18 Å from data with a physical Nyquist limit of 4.78 Å. These results support a practical acquisition strategy: a lower magnification can be selected to increase the FOV, capturing more cellular context and more particles per tilt series. Processing data at the super-resolution pixel size can recover information beyond the physical Nyquist limit if the chosen sampling limits resolution.

## Results

### *In situ* cryo-ET reveals putative virus particles in *Drosophila melanogaster* larvae

We performed serial lift-out on high-pressure frozen third-instar *D. melanogaster* larvae to prepare lamellae for subsequent cryo-ET analysis. We targeted the interface between the cuticle and the apical membrane using sample autofluorescence together with the fluorescence from mNeonGreen-tagged chitin synthase (Fig. S2; see Methods) ^31,32^. Four lamellae were prepared during a single 24-hour lift-out session (Fig. S2). We collected tilt series at a nominal magnification of 53,000×, corresponding to a physical pixel size of 2.39 Å/px and a super-resolution pixel size of 1.195 Å/px. Each tilt image covered approximately 0.96 µm^2^. We collected 32 tilt series from the four lamellae, of which 28 produced tomograms of sufficient quality for analysis.

Tomograms reconstructed at 14.34 Å/px revealed electron-dense, spherical particles at the interface between the cuticle and the apical membrane (Fig. 1a, Fig. S2, and Supplementary Video 1). These particles were approximately 30 nm in diameter and occurred both intracellularly and within the cuticle (Fig. 1a). They appeared as isolated particles in some tomograms and as tightly packed, regular lattice arrays in others (Fig. 1a, Supplementary Video 1). Their morphology, size, and tendency to form ordered arrays led us to hypothesise that they were virus particles, consistent with previously described cryo-ET observations of viruses *in situ* ^33–39^. The putative viral infection was serendipitous, and no similar particles were observed in other biological replicates of the third-instar *D. melanogaster* larvae we analysed. The total number of available tomograms containing the putative virus particles was therefore limited. Of the 28 reconstructed tomograms, 20 contained these virus particles and were used for subsequent STA.

**Figure 1.**
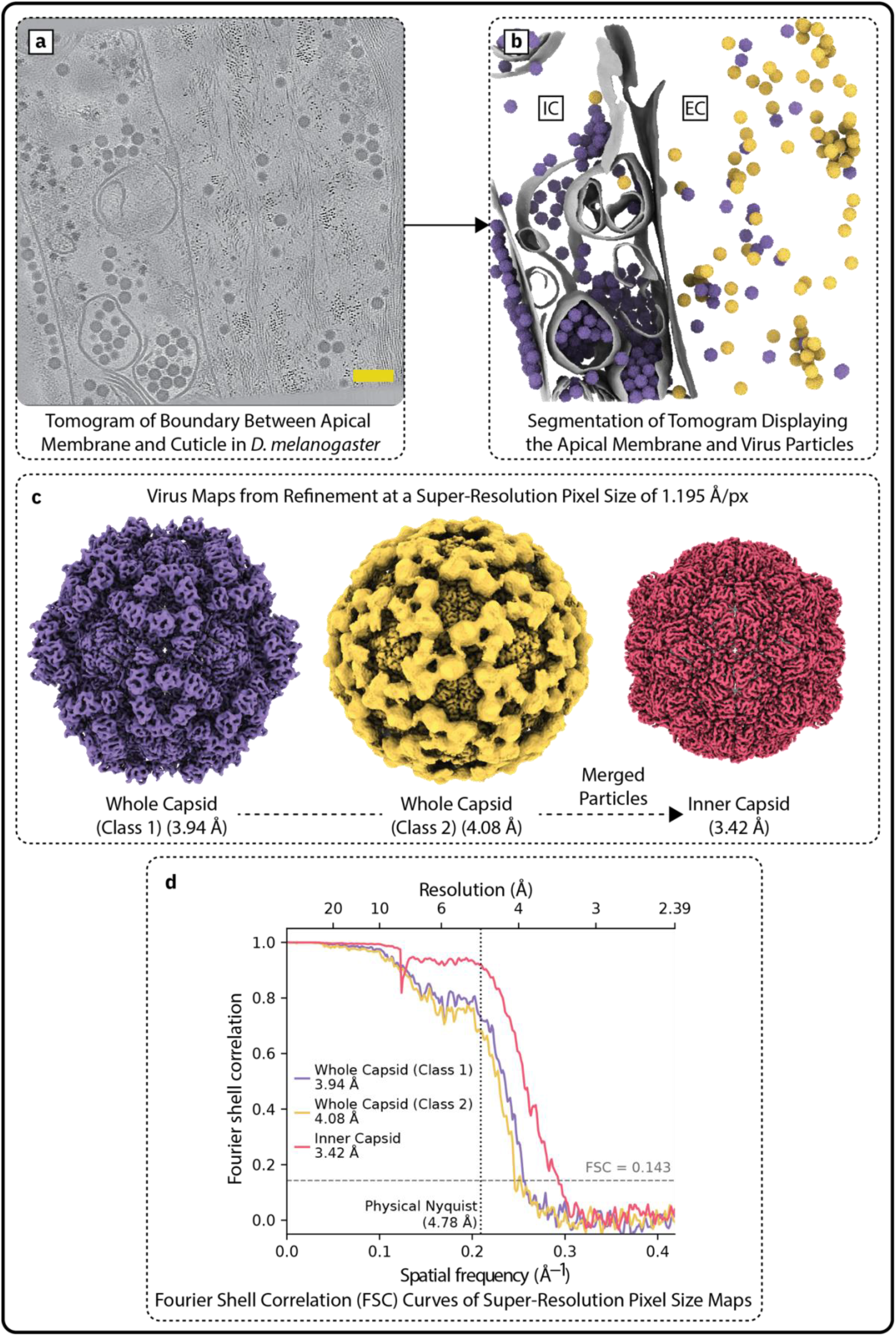
Super-resolution STA resolves virus capsids in *Drosophila* larval tissue beyond the physical Nyquist limit. (a) Slice of an IsoNet2 denoised tomogram at 14.34 Å/px at the interface between the apical membrane and the cuticle in *D. melanogaster*. Scale bar = 100 nm. (b) Segmentation of the tomogram in *a*. Membranes are coloured grey, with the apical membrane coloured dark grey. Virus particles, after STA and classification, were back-plotted into the tomogram. Class 1 is coloured purple and Class 2 is coloured yellow. The intracellular space is found to the left of the apical membrane and is labelled IC. The extracellular space, here surrounded by the chitinous cuticle, is found to the right of the apical membrane and is labelled EC. (c) Virus maps produced by STA at super-resolution pixel size (1.195 Å/px). All resolutions are reported using a Fourier shell correlation (FSC) threshold of 0.143. Colours for Class 1 and Class 2 are the same as in *b.* The inner capsid reconstruction (Class 1 and 2 particles combined) is coloured red. All particles are shown in the same orientation. Maps shown were denoised by M for display. (d) FSC curves for the reconstructions in *c*, with the 0.143 threshold and the physical Nyquist frequency (4.78 Å) marked.

### Super-resolution STA enables *de novo* model building and identification of Drosophila A virus

We performed STA on the putative viral particles from all 20 tomograms in which they were detected. Initial alignments and 3D classifications separated the particles into two classes, both consisting of small, non-enveloped particles with icosahedral capsids (Fig. S3a). The two classes were refined separately with imposed icosahedral symmetry. At the physical pixel size of 2.39 Å/px, the Fourier shell correlation (FSC) curves for both reconstructions remained above the 0.143 threshold at the physical Nyquist limit of 4.78 Å, indicating that their resolution was limited by sampling (Fig. S3b). The inner capsids of the two classes were visually indistinguishable (Fig. S3c) and had a real-space correlation coefficient of 0.88 within a masked volume containing only the inner capsid. In contrast, the structures of the outer capsid differed markedly (Fig. S3a). Class 1 exhibited a well-ordered outer capsid with icosahedral symmetry, whereas the outer capsid of Class 2 was poorly resolved, irrespective of whether symmetry was applied (Fig. S3a). The two classes therefore appeared to represent the same virus with different outer capsid organisations. Upon back-plotting the positions of each particle into the tomograms, we found that 63% of Class 1 particles were intracellular, whereas 91% of Class 2 particles were located within the cuticle (Fig. 1b, Supplementary Video 2).

To identify the virus, we sought to build a protein model *de novo* and use the resulting sequence to search the BLAST database. The maps obtained at the physical pixel size were not sufficiently resolved for reliable, automated *de novo* model building with tools such as ModelAngelo ^40^, which typically require resolutions better than 4 Å. Because both reconstructions had reached the physical Nyquist limit, we re-rendered the original EER movies at the super-resolution pixel size of 1.195 Å/px and repeated a limited STA processing workflow (see Methods). This improved the resolution of Class 1 (3,513 particles) to 3.94 Å and of Class 2 (2,342 particles) to 4.08 Å (FSC = 0.143, whole capsid mask; Fig. 1c,d). Since the inner capsids were indistinguishable, we combined all 5,855 particles and refined them using an inner capsid mask. The resulting reconstruction reached 3.42 Å resolution, well beyond the physical Nyquist limit of 4.78 Å (FSC = 0.143, inner capsid mask; Fig. 1c,d).

A Rosenthal-Henderson plot ^41^ of the inner capsid reconstruction indicated that approximately 53 virus particles (∼3,180 asymmetric units) were required to exceed the physical Nyquist limit (Fig. S4). The particle subsets were reconstructed using alignment and deformation parameters obtained from the full dataset and were not refined independently. This estimate therefore describes the particle number required at the alignment accuracy of the final refinement, rather than the number of particles required for independent particle alignment.

The resolution improvement achieved by this approach was also apparent in the reconstructed maps. Refinement at the super-resolution pixel size produced more clearly resolved features than refinement at the physical pixel size (Fig. 2). Simply upsampling the physical-pixel reconstruction to 1.195 Å/px without further refinement did not reproduce these features (Fig. 2). The improvement therefore resulted from the recovery and refinement of signal beyond the physical Nyquist limit rather than from interpolation.

**Figure 2.**
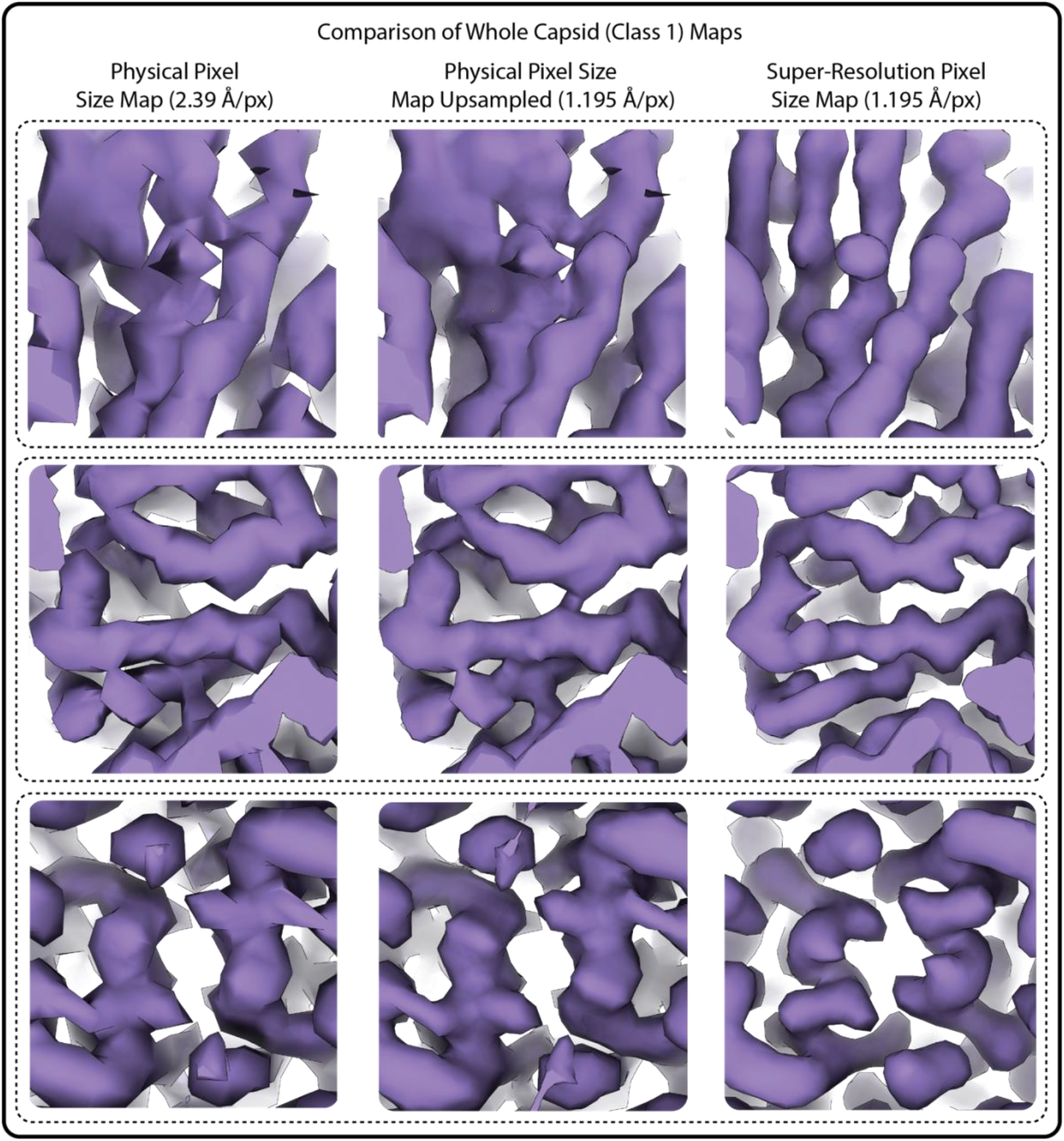
Super-resolution processing improves capsid density beyond simple map upsampling. Three representative regions of the Class 1 capsid reconstruction are shown under three processing conditions. Left: reconstruction refined at a physical pixel size of 2.39 Å/px. Middle: the same reconstruction resampled in ChimeraX ^42^ to 1.195 Å/px without further refinement. Right: reconstruction obtained by re-rendering the EER data at a super-resolution pixel size of 1.195 Å/px and repeating refinement. Maps shown were denoised by M for display.

The 3.42 Å inner capsid map supported automated model building using ModelAngelo without a prior sequence (Fig. S5) ^40^. The most complete chain contained 169 residues. Its predicted sequence returned the capsid protein of Drosophila A virus (DAV) as the highest scoring BLAST match with 63% identity and an E-value of 3×10⁻⁶⁰. The sequence was not expected to be fully accurate because ModelAngelo could not assign every side chain correctly without a sequence prior.

To test whether the sequence assignment was consistent with the structure, we truncated an AlphaFold ^43^ model of the DAV capsid protein to only include the inner capsid region and rigid-body fitted it into the map. The predicted structure agreed with the map, with a main chain map-model correlation coefficient (CC) of 0.69 (Fig. S5). We then repeated automated model building using ModelAngelo ^40^ and the DAV capsid protein sequence as a prior, fitted the model into our map using Coot, and refined it in Phenix (Fig. 3a,b) ^44,45^. Most side-chain density was resolved and the final model agreed closely with the map (Fig. 3b), with a side-chain and main chain map-model CC of 0.83 and 0.85, respectively. The inner capsid was found to form a T=3 lattice, consistent with the previously reported 8 Å *in vitro* reconstruction of DAV (Fig. 3a) ^46^. On this basis, we identified the particles as Drosophila A virus. Super-resolution processing thus enabled *de novo* atomic model building and sequence assignment without prior sequence information, from 5,855 particles in 20 tomograms.

**Figure 3.**
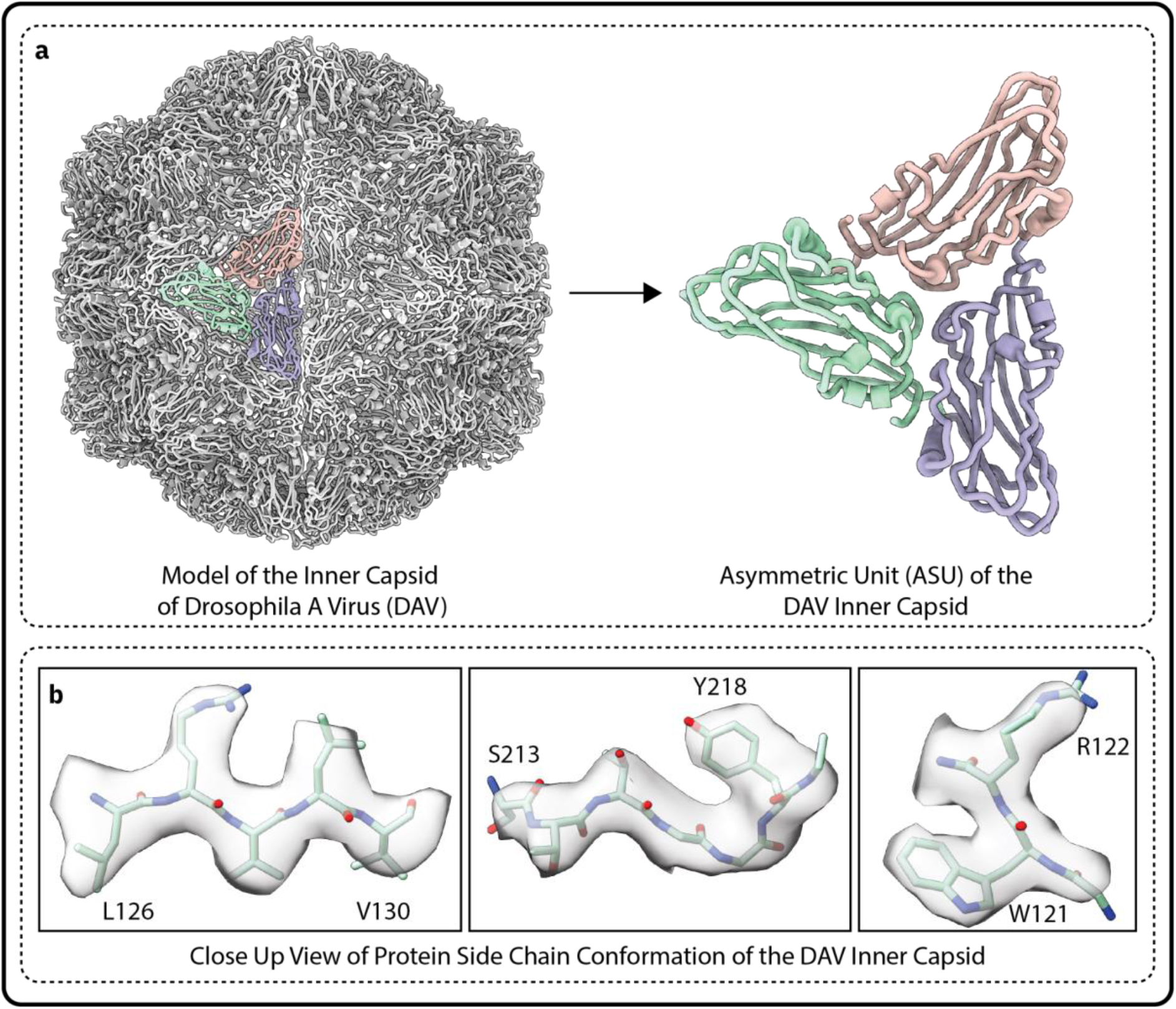
*De novo* model building identifies Drosophila A virus (DAV). (a) Refined atomic model of the DAV inner capsid. One asymmetric unit is highlighted within the T=3 capsid; its three sequence-identical, quasi-equivalent subunits are shown in green, pink and blue, while all other subunits are grey. An enlarged view of the highlighted asymmetric unit is shown on the right. (b) Representative close-up views of the density and fitted model. Resolved side-chain features agree with the DAV capsid sequence and support the sequence assignment. The map was filtered according to its estimated resolution and sharpened using M for display.

### Super-resolution STA resolves *in situ* ribosomes beyond the physical Nyquist limit

Icosahedral viruses are particularly favourable targets for STA. Their symmetry provides 60 copies of the asymmetric unit per virus particle, while their large, rigid capsids can be aligned accurately despite the low signal-to-noise ratio of cryo-ET data. High-resolution reconstructions of icosahedral viruses can therefore be obtained from relatively few particles^36^. In our Drosophila A virus dataset, the Rosenthal-Henderson plot indicated that approximately 53 virus particles were sufficient to exceed the physical Nyquist limit (Fig. S4). Most targets studied by *in situ* STA lack such symmetry, and for asymmetric complexes each subtomogram contributes a single copy of the structure to the average. Recovering information beyond the physical Nyquist limit for an asymmetric target is therefore expected to require substantially more particles. To test whether super-resolution processing also benefits an asymmetric target in a typical *in situ* STA dataset, we analysed cytoplasmic ribosomes in *S. cerevisiae* at the super-resolution pixel size.

We plunge-froze *S. cerevisiae* cells and prepared lamellae by cryo-focused ion beam (FIB) milling (see Methods). As ribosomes are abundant throughout the cytoplasm, fluorescence-guided targeting was not required. We prepared 25 lamellae within a single 24-hour session and collected tilt series using the same imaging conditions used for the *D. melanogaster* dataset. This corresponded to a physical pixel size of 2.39 Å/px, a super-resolution pixel size of 1.195 Å/px, and an imaged area of approximately 0.96 µm² per tilt image. We collected 349 tilt series from 25 lamellae, of which 319 produced tomograms of sufficient quality for analysis.

The EER movies were rendered at the super-resolution pixel size, and the tomograms were reconstructed as described for the DAV dataset (see Methods). Ribosomes were identified by template matching using PyTOM ^47^, yielding 190,883 particles from 310 tomograms. Following refinement in M, the consensus ribosome reached 3.18 Å (FSC = 0.143), well beyond the physical Nyquist limit of 4.78 Å (Fig. 4a,c). Local resolution reached a maximum of 2.78 Å in the core of the 60S subunit, whereas more flexible regions, including parts of the 40S subunit, were resolved at lower resolution (Fig. 4b). The map contained features consistent with the reported resolution, including resolved bulky side chains (Fig. 4d). These results show that information beyond the physical Nyquist limit can be reproducibly recovered also for asymmetric *in situ* targets.

**Figure 4.**
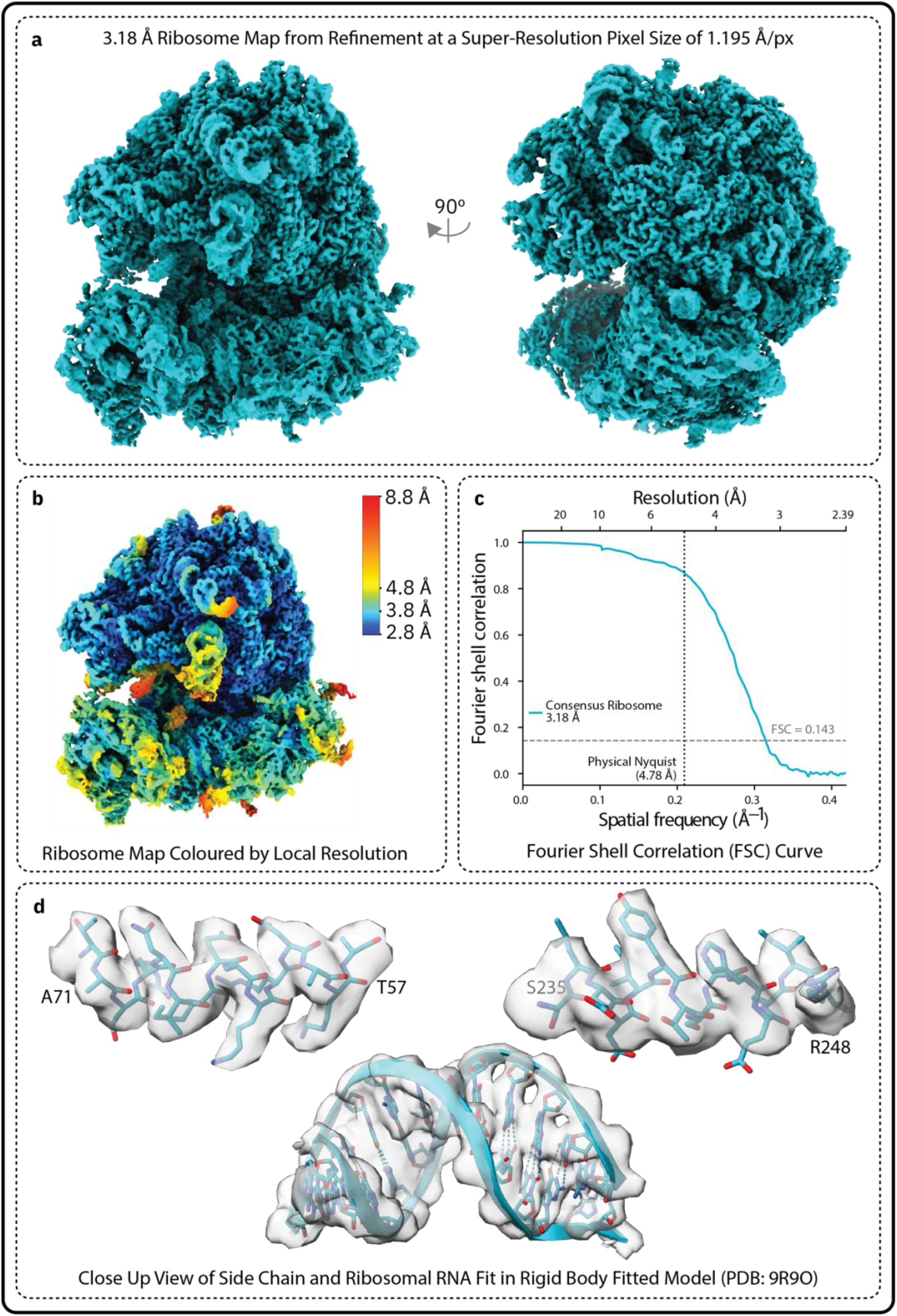
Super-resolution STA resolves the *S. cerevisiae* ribosome beyond the physical Nyquist limit. (a) Two views of the consensus ribosome map reconstructed from 190,883 particles at the super-resolution pixel size of 1.195 Å/px. The global resolution reached 3.18 Å, as measured at an FSC threshold of 0.143. (b) Consensus map coloured by local resolution estimated in M. Resolution reached 2.78 Å in the core of the 60S subunit. (c) FSC curve for consensus reconstruction with the 0.143 threshold and the physical Nyquist frequency (4.78 Å) marked. (d) A previously determined model of the *S. cerevisiae* ribosome (PDB 9R9O) rigid-body fitted into the map. Enlarged views show resolved protein side chains and ribosomal RNA features. Maps shown in *a, b,* and *d* were denoised by M for display.

## Discussion

This study shows that processing Falcon 4i EER data at a super-resolution pixel size recovers structural information beyond the physical Nyquist limit for *in situ* structure determination by cryo-ET. Reconstructions of both the Drosophila A virus inner capsid and the *Saccharomyces cerevisiae* ribosome reached resolutions better than 3.5 Å, well beyond the Nyquist limit of the acquired data. This was achieved despite conditions that lower the signal-to-noise ratio: the *Drosophila* lamellae were 100–270 nm thick, compared with <60 nm ice in typical SPA specimens, contained 20% dextran, which lowers contrast ^48^, and carried ion-beam damage from FIB milling ^49^. A previous attempt at super-resolution STA on purified apoferritin reported FSC resolutions beyond the physical Nyquist limit that were not supported by map features, and concluded that it gave no benefit ^30^. Here, resolved side-chain density consistent with resolution estimates and a coherent *de novo* sequence assignment provide evidence independent of the FSC, which neither interpolation nor aliasing can produce. The extent of improvement will depend on the specimen and target. However, the two datasets differ substantially in particle size and symmetry, and in specimen and lamella preparation. Our results therefore show that recovery beyond the physical Nyquist limit is not restricted to large, highly symmetric viral particles.

The practical implications of these results concern the choice of pixel size when establishing data-acquisition routines. EER acquisition makes it possible to use a lower magnification to increase the field of view and the number of particles recorded per tomogram, while retaining finer effective sampling through super-resolution rendering. Finer sampling alone, however, does not guarantee that high-resolution information can be recovered. The DQE decreases at spatial frequencies beyond the physical Nyquist limit, reducing the signal contributed by each electron relative to noise ^26,27,50^. In SPA, Feathers *et al.* showed that the additional particles captured within a larger FOV can compensate for the reduced DQE at high spatial frequencies ^29^. This strategy has seen limited use in SPA because particles are generally abundant across the grid. It is therefore usually straightforward to collect additional micrographs at a physical pixel size compatible with the target resolution, without incurring the loss of DQE associated with super-resolution imaging.

*In situ* cryo-ET imposes an acquisition constraint absent from SPA: tilt series positions must be spaced by the area illuminated throughout the tilt series, not by the area recorded at zero tilt. Perpendicular to the tilt axis, the illuminated area grows with tilt angle, reaching twice the beam footprint at 60°, so adjacent regions cannot be recorded without pre-exposing them ^51,52^. A particle cluster extending beyond the FOV used for high-resolution acquisition therefore cannot be completed by a neighbouring tilt series. A larger FOV can capture the entire cluster in a single tilt series, increasing particle yield per lamella. Lower-magnification acquisition combined with super-resolution processing is therefore most valuable for sparse, clustered targets. A larger FOV also tolerates error in positioning tilt series on targets that are difficult to locate in lamella overviews, such as fluorescence-targeted regions in lift-out lamellae. More generally, a larger FOV preserves additional cellular context and allows a given area to be covered with fewer tilt series, reducing the microscope time required to image the specimen. These advantages may justify acquisition at lower magnification even when the target is uniformly distributed and the total particle yield per lamella is expected to remain largely independent of magnification. Other strategies address the same constraints. Tilt schemes that tolerate controlled beam overlap allow denser spacing of positions ^53,54^, and low-magnification context tomograms can guide high-magnification acquisition ^55–57^. The latter splits context and structure into two datasets, whereas super-resolution processing provides both from a single tomogram, at the cost of reduced DQE beyond the physical Nyquist limit.

A pilot dataset can help guide the choice of pixel size before committing to a large-scale acquisition for high-resolution *in situ* structure determination. Processing can initially be performed at the physical pixel size to contain computational and storage costs. If the resulting reconstruction approaches the physical Nyquist limit, the original EER files can be re-rendered at the super-resolution pixel size and the relevant processing and refinement steps repeated. Particle coordinates and other information obtained during the initial processing can be reused, as demonstrated for the DAV dataset. Super-resolution processing therefore provides a practical route for revisiting existing datasets that prove to be limited by sampling, provided that the original EER data have been retained.

The physical Nyquist limit remains an important consideration when planning data acquisition, but it does not necessarily constitute an absolute resolution ceiling for *in situ* STA. When sufficient particles are available, information beyond this limit can be recovered. The most appropriate acquisition strategy will depend on the specimen, the distribution of the target and the experimental objective. Nevertheless, EER acquisition makes the choice of pixel size less rigid; magnification can be selected with greater consideration for field of view, cellular context and particle yield, while preserving the option to process the data at super-resolution if physical sampling becomes limiting.

## Supporting information

supplementary_scripts

## Acknowledgments

We thank members of the Mattei lab, particularly Higor Rosa, and the cryo-EM community at EMBL for useful discussions and technical support. We would also like to thank Simon Fromm and Jenny Sachweh for critical readings of the manuscript. EP and VD were funded by the Deutsche Forschungsgemeinschaft (DFG, German Research Foundation) – SPP2416 project number 525894472. EP was funded by the EMBL Interdisciplinary Postdoc Program (EIPOD). SB was funded by the Deutsche Forschungsgemeinschaft (DFG, German Research Foundation) – SPP2453 project number 541626282. SM was funded by the European Molecular Biology Laboratory. We acknowledge the access and services provided by the Imaging Centre at the European Molecular Biology Laboratory (EMBL IC), generously supported by the Boehringer Ingelheim Foundation. During the preparation of this work, the authors used Claude (Anthropic) and ChatGPT (OpenAI) to assist with drafting and revising prose, literature search support, and writing code. After using these tools, the authors reviewed and edited the content as needed and take full responsibility for the content of the publication.

## Author Contributions

Conceptualisation, EP; *Drosophila* culture and sample preparation, VD; serial lift-out, EP; whole-cell FIB milling, SB; cryo-ET data collection, EP and SB; cryo-ET and STA data processing, EP and SB; manuscript writing, EP; manuscript revisions, EP, SM; funding acquisition, SM.

## Data Availability

The STA maps and model coordinates have been deposited in the Electron Microscopy Data Bank (EMDB) and in the Protein Data Bank (PDB). Drosophila A Virus (Whole Capsid) (Class 1): EMD-XXX. Drosophila A Virus (Whole Capsid) (Class 2): EMD-XXX. Drosophila A Virus (Inner Capsid): EMD-XXX, PDB XXXX. *S. cerevisiae* Consensus Ribosome: EMD-XXXX.

## Methods

### *Drosophila melanogaster* culture and high-pressure freezing

*Drosophila melanogaster* expressing mNeonGreen-tagged Chitin Synthase 1 (w[*]; TI{TI}kkv[mNG.DsRed-]; RRID:BDSC_84992) were raised on standard fly food at 25 °C. Larvae were staged according to Sui and Dahmann ^58^, and mid-third-instar larvae were collected at 80–85 hours after egg laying. For high-pressure freezing, individual larvae were dissected in Grace’s Insect Medium, prepared according to Sui and Dahmann ^58^. Dissection involved removing the anterior and posterior regions, gut, and fat body. The dissected larvae were placed dorsal side up in the 100-µm well of a 3-mm Type A sample carrier (Leica Microsystems) containing Grace’s Insect Medium supplemented with 20% (w/v) dextran as a cryoprotectant. High-pressure freezing was performed using an EM ICE high-pressure freezer (Leica Microsystems).

### Fluorescence microscopy of high-pressure frozen carriers

Fluorescence microscopy of high-pressure frozen carriers was performed using a Zeiss LSM 900 Airyscan 2 microscope with a cryo-stage (Linkam). Images were acquired using a 10× objective in confocal mode. A 488-nm laser was used to acquire images of mNeonGreen-tagged chitin synthase fluorescence. Illumination and detection were controlled by ZEN Blue software which was also used to stitch tile images into montages. Most of the signal arose from specimen autofluorescence, which was used to map the morphology of the larva within the carrier. Chitin synthase localises to plaques on the tips of the apical membrane undulae at the interface of the apical membrane and the chitinous cuticle, which appear in the images as fluorescent puncta ^31,32^. Carriers containing regions which displayed numerous fluorescent puncta were selected for serial lift-out to target the interface between the apical membrane and the cuticle.

### Serial lift-out of *D. melanogaster* larvae

Serial lift-out was performed on a single high-pressure frozen larva, similar to a previously described workflow ^12^. A high-pressure frozen carrier selected for serial lift-out by fluorescence microscopy was loaded into a 35° pre-tilted Aquilos shuttle (Thermo Fisher Scientific) in cryo-conditions. A clipped UltrAuFoil, R2/2 200 mesh grid (Quantifoil) was loaded into the other shuttle slot for use as a receiver grid. The shuttle was loaded into an Aquilos 2 focused ion beam-scanning electron microscope (FIB-SEM) with an integrated fluorescence microscope and an EasyLift needle system (Thermo Fisher Scientific). The region intended for lift-out, which was selected during previous cryo-fluorescence microscopy, was located within the intact high-pressure frozen carrier using the integrated fluorescence microscope. The sample was then protected from beam damage by applying a metal-organic platinum layer using the gas injection system (GIS). Trenches surrounding the region of interest were milled using a current of 16 nA, creating a block of ice containing the region of interest. An undercut of the block of ice was then performed using a current of 3 nA. A copper block on the needle was attached to the block of ice using redeposition. The ice block was freed from the carrier by milling the last remaining attachment point of the block to the carrier.

The ice block was then moved to the receiver grid using the needle. The ice block was sequentially milled using the FIB to deposit a series of 3.5 µm thick slices onto the foil of the receiver grid with a current of 1 nA, according to the SOLIST method ^21^. Another protective metal-organic platinum layer was applied using the GIS. Slices were then thinned using decreasing milling currents depending on sample thickness: 1 nA to 3 µm, 0.5 nA to 2 µm, 0.3 nA to 1 µm, 50 pA to 500 nm, 30 pA to 180 nm. The resulting lamellae were 12 µm wide and were then polished using 30 pA at a 0.5° over-tilt. The lamellae were sputter coated at 10 mA for 3 seconds to reduce charging during cryo-ET ^59^. In total, 4 lamellae were generated in a single lift-out session over 24 hours.

### Cryo-electron tomography data collection of *D. melanogaster* larvae

The receiver grid from serial lift-out was loaded onto a Titan Krios G4 operated at 300 kV and was equipped with a Selectris X energy filter and a Falcon 4i DED (Thermo Fisher Scientific). Images were acquired using SerialEM v4.2.5 set up using the LowDoseWizard ^60,61^. Prior to tilt series acquisition, the entire area of each lamella was imaged at a pixel size of 28.78 Å/px and the images were stitched together to create an overview of the sample. These overviews were used to select the acquisition point for each tilt series. Tilt series were acquired with PACE-tomo ^52^ at a nominal magnification of 53,000×, corresponding to a physical pixel size of 2.39 Å/px, in nanoprobe mode with a 50 µm C2 aperture and a 70 µm objective aperture inserted. The energy filter slit was inserted at a width of 10 eV. Tilt series were acquired using a dose-symmetric tilt scheme ^62^ with a 3° step over a total range of −51 to +51°, relative to the lamella pre-tilt, yielding 35 tilt images per tilt series. The nominal defocus was set between −3 and −6 µm. Each tilt image was exposed for 1.14 seconds at an electron dose of ∼3.65 e⁻/Å², giving a total electron dose of ∼128 e⁻/Å² per tilt series. Movies were recorded in EER format ^25^. A total of 32 tilt series were collected from 4 lamellae.

### Tomogram reconstruction of *D. melanogaster* larvae

Tilt series were processed using the WarpTools and M pipeline for Linux (Warp v2.0.0dev36) ^1,63^ with all steps being carried out using this software, unless stated otherwise. EER movies were converted to .tif format at the physical pixel size (2.39 Å/px) using relion_convert_to_tiff (RELION v5) ^2^, grouping every 25 EER frames to yield 14 fractions per tilt image. Motion correction and contrast transfer function (CTF) estimation were carried out on a 1×1×14 motion grid and a 2×2×1 CTF grid, fitting the CTF between 40 and 8 Å. Tilt series stacks were then generated. These stacks were manually inspected and poor-quality images were removed from further processing using custom scripts which modified the .mdoc files. AreTomo2 (v1.0.0) ^64^ was used for tilt series alignment at 14.34 Å/px. Preliminary tomograms were used to estimate the ice thickness of each tomogram, and these thickness values were used to set the -AlignZ value for each tilt series during a second round of AreTomo2 alignment. The defocus hand was then estimated, and the tomograms flipped to ensure correct tomogram defocus handedness. 3D-CTF corrected tomograms were generated at a pixel size of 14.34 Å/px. A total of 28 tomograms were of a sufficient quality to interpret and analyse.

### Tomogram denoising and segmentation of *D. melanogaster* larvae

Tomograms were regenerated at 14.34 Å/px from odd and even frame half-sets and were denoised and missing-wedge corrected using IsoNet2 ^65^. A noise2noise denoising network was first trained and applied (CTF mode ‘network’, phase-flipped input). The denoised tomograms were then used to generate masks and to train an IsoNet2 noise2noise refinement network with a missing-wedge weight of 0, which was used to predict the final tomograms. Denoised tomograms were solely used for visualisation and membrane segmentation and were not used in any STA steps.

Membranes were segmented on the IsoNet2 denoised tomogram using MemBrain-seg ^66^. The final positions and orientations of Class 1 and Class 2 virus particles were read from the .star files produced at the end of STA (see below) and mapped back into the segmentation using ArtiaX ^67^. Segmentations were visualised within ChimeraX ^42^.

### STA of virus particles at the physical pixel size

Particles were extracted as 3D subtomograms by WarpTools and STA was carried out in RELION (v4) ^68^, unless stated otherwise. To generate an initial reference, 1,458 virus particles were manually picked in 4 tomograms. Particles were extracted at 19.12 Å/px and an initial model was generated *de novo* without imposed symmetry. Particles were re-extracted at 9.56 Å/px. 3D classification (T = 1, K = 4) with alignment and without imposed symmetry separated two classes of virus particle. Particles from the class which refined to the highest resolution were re-extracted and refined at 4.78 Å/px. The resulting map was used as a template for template matching.

Template matching was carried out using PyTOM ^47^ on 3D-CTF corrected tomograms at 14.34 Å/px, using a template box of 40 voxels, a spherical mask, and a particle diameter of 430 Å. A total of 10,691 particles were picked. Particles were extracted at 9.56 Å/px and subjected to 3D classification with alignment without imposed symmetry (T = 1, K = 6), using the map from the manually picked particles as a reference. This classification separated Class 1 and Class 2. Each class was refined separately with icosahedral symmetry (I2) at 9.56 Å/px, then at 4.78 Å/px, and finally at 2.39 Å/px, reaching resolutions of 9.1 Å (Class 1) and 9.8 Å (Class 2). To resolve the outer capsid, Class 2 was further classified and refined with and without imposed symmetry, but no class showed improved outer capsid density.

Particles were then refined in M. Initially, particle poses (refine_particles) and tilt series alignment (refine_imagewarp 1×1) were refined, followed by per-tilt defocus once the resolution reached 8 Å (ctf_defocus, initially with ctf_defocusexhaustive). Over subsequent rounds in M, the image warp grid was increased to 8×8, and volume warp (refine_volumewarp 8×8×4×35), third-order Zernike aberrations (ctf_zernike3), and per-particle and per-frame weights (EstimateWeights) were introduced. Both classes reached 4.81 Å, which coincided with the physical Nyquist limit of 4.78 Å, and could not improve further.

### Reprocessing of viruses at the super-resolution pixel size

The data were re-processed at the super-resolution pixel size from the original EER movies of the 20 tomograms containing virus particles used for STA. Movies were rendered at 1.195 Å/px using relion_convert_to_tiff with an EER upsampling factor of 2, grouping every 25 EER frames as before. The warp_frameseries.settings and warp_tiltseries.settings files were generated (create_settings) using the new super-resolution pixel size. The tomogram dimensions, which are given in pixels, were doubled to maintain the same tomogram size. Motion correction and CTF estimation were repeated in WarpTools using identical parameters. The tilt image exclusions from physical pixel size processing were retained by reusing the modified .mdoc files. Tilt series alignment was not repeated. Instead, the tilt series alignments from AreTomo2 were imported without modification by soft-linking to the tiltstack directory in the physical pixel size processing directory, and importing the alignments (ts_import_alignments; at an alignment pixel size of 14.34 Å/px as per physical pixel size processing). The defocus hand was checked and set, and tomograms were reconstructed at 14.34 Å/px.

Particle picking was not repeated. Instead, the coordinates from physical pixel size processing were used. As a precaution against minor changes in particle positions due to repeated motion correction, the particle coordinates and orientations from the final refinements of Class 1 and Class 2 at 2.39 Å/px were used to extract particles at 9.56 Å/px. Each class was re-refined in RELION at 9.56 Å/px and then at 4.78 Å/px with I2 symmetry. Duplicate particles were removed. New M species were created from these refinements with the particles resampled to 1.195 Å/px, an initial low-pass filter of 10 Å, and I2 symmetry. M refinement was carried out as described above, progressively increasing the image warp grid from 1×1 to 8×8 and the volume warp grid from 4×4×2×35 to 8×8×4×35, and refining stage angles (refine_stageangles), per-tilt defocus, and third-order Zernike aberrations. Per-particle, per-frame and anisotropic weights were estimated with EstimateWeights. A final round of refinement included per-tilt movie alignment (refine_tiltmovies). Class 1 reached 3.94 Å from 3,513 particles in 20 tomograms and Class 2 reached 4.08 Å from 2,342 particles in 19 tomograms. Resolutions were determined by relion_postprocess using the M half-maps and a mask covering the whole capsid (extended by 2 pixels with a soft edge of 8 pixels), with an FSC threshold of 0.143.

An inner capsid map was generated by combining particles from both classes and refining the map with an inner capsid mask. The mask covering only the inner capsid was generated by segmenting the Class 2 map in ChimeraX using Segger, and a soft-edged mask (8 pixels) was created in RELION. To refine the particles, a new M species was created from the .star file containing both classes, and using the Class 2 half-maps as a reference with an initial low-pass filter of 3.9 Å with I2 symmetry. The combined species was refined in M using image warp (8×8), particle pose refinement, per-tilt defocus or third-order Zernike aberrations, and volume warp (8×8×4×35), with per-particle, per-frame and anisotropic weights re-estimated with EstimateWeights. A final round of refinement used a volume warp grid of 10×10×6×35 and additionally refined fifth-order Zernike aberrations and anisotropic magnification. The inner capsid reconstruction reached 3.42 Å from 5,855 particles in 20 tomograms (FSC = 0.143, inner capsid mask). Metrics for the data collection and processing of all maps are summarised in Supplementary Table 1 and 2.

### *De novo* model building and identification of Drosophila A virus

Automated model building was carried out without a sequence prior using ModelAngelo (v1.0.9) ^40^ on the masked, sharpened 3.42 Å inner capsid map. The sequence of the most complete chain (169 residues) was searched using BLASTp ^69^. The highest-scoring hit was the capsid protein of Drosophila A virus (DAV) (GenBank AWY11071.1; 63.3% identity over 169 residues, 99% query coverage, E-value = 3×10⁻⁶⁰).

To validate this assignment, the structure of the DAV capsid protein was predicted using the AlphaFold 3 server ^70^. The predicted model was truncated to the inner capsid and rigid-body fitted into the inner capsid map using fitmap in ChimeraX ^42^. The pLDDT values of the predicted model were converted to B-factors using phenix.process_predicted_model ^71^, and the map-model correlation was calculated with phenix.validation_cryoem using the half-maps (main chain CC = 0.69) ^72^.

### Model building and refinement

ModelAngelo ^40^ was run again using the DAV capsid protein sequence as a prior on the inner capsid map. One asymmetric unit, comprising three chains, and its immediate neighbours were selected from the resulting model. Each chain was manually rebuilt in Coot (v0.9.7) ^45^ into the sharpened inner capsid map, and the three chains of the asymmetric unit were trimmed to cover the same residues (residues 69–240). The icosahedral symmetry operators were determined from the map using phenix.map_symmetry, and the asymmetric unit together with its immediate neighbours was generated with phenix.apply_ncs. The model was refined using phenix.real_space_refine (Phenix v2.1) ^44^ at a resolution of 3.5 Å. Neighbouring chains were regenerated from the asymmetric unit by symmetry before each round of refinement, and from the third round onwards were restrained to their starting coordinates during Phenix refinement (reference coordinate restraints, σ = 0.1 Å). Refinement in Phenix and manual correction in Coot, guided by phenix.validation_cryoem, per-residue map-model correlations (phenix.map_correlations), were iterated for four rounds. The final asymmetric unit was validated with phenix.validation_cryoem using the half-maps (CC_mask = 0.87; main chain CC = 0.85; side chain CC = 0.83) and MolProbity ^73^. The complete inner capsid was generated by applying icosahedral symmetry to the asymmetric unit. Model statistics are summarised in Supplementary Table 3 and 4.

### Comparison of Class 1 and Class 2

To compare the inner capsids of Class 1 and Class 2, a mask covering only the inner capsid was generated from a 10 Å simulated map (molmap) of the refined inner capsid model in ChimeraX ^42^. The real-space correlation coefficient between the Class 1 and Class 2 half-maps at the physical pixel size was calculated within this mask using a custom Python script (Supplementary Script 1). To determine the location of each class, Class 1 and Class 2 particles were mapped back into each tomogram in ArtiaX, and particles located within the cuticle were manually counted for each class.

### *Saccharomyces cerevisiae* culture and plunge-freezing

*S. cerevisiae* (BY4741 strain) samples were grown in YPGly medium at 30 °C to an OD₆₀₀ of 1. R2/2 200 Mesh Cu (Quantifoil) grids were plasma-cleaned for 30 s in 95% argon/5% oxygen using a 1070 plasma cleaner (Fischione). 3 µL of cells were applied to the grids, back-blotted for 1.2 seconds, and plunge-frozen in liquid ethane using a Leica EM GP2 plunger at 25 °C and 90% humidity.

### Cryo-FIB milling of *S. cerevisiae*

Plunge-frozen grids containing *S. cerevisiae* were clipped into Autogrids and loaded into a 35° pre-tilted Aquilos shuttle (Thermo Fisher Scientific). The shuttle was loaded under cryo-conditions into the same Aquilos 2 FIB-SEM (Thermo Fisher Scientific) used for serial lift-out. The grid was sputter coated before a protective metal-organic platinum layer was applied using the GIS, and the grid was then sputter coated again. Lamellae were milled using AutoTEM software (Thermo Fisher Scientific) with decreasing milling currents depending on sample thickness: 1 nA to 4 µm, 0.5 nA to 2 µm, 0.3 nA to 1 µm, 50 pA to 500 nm, 30 pA to 200 nm. The resulting lamellae were 15 µm wide and were then polished using 30 pA at a 0.5° over-tilt. Lamellae were sputter coated at 10 mA for 4 seconds. A total of 25 lamellae were generated in 24 hours. As ribosomes are abundant throughout the cytoplasm, no targeting by fluorescence was required.

### Cryo-electron tomography data collection of *S. cerevisiae*

The grid from *S. cerevisiae* cryo-FIB milling was loaded onto the same Titan Krios G4 used for the *D. melanogaster* data. The microscope was operated at 300 kV and was equipped with a Selectris X energy filter and a Falcon 4i DED (Thermo Fisher Scientific). Images were acquired using SerialEM v4.3.0beta set up using the LowDoseWizard ^60,61^. Tilt series were acquired on yeast cells, avoiding regions containing no or limited numbers of ribosomes, such as the nucleus. Tilt series were acquired with PACE-tomo ^52^ at a nominal magnification of 53,000×, corresponding to a physical pixel size of 2.39 Å/px, in nanoprobe mode with a 50 µm C2 aperture and a 70 µm objective aperture inserted. The energy filter slit was inserted at a width of 10 eV. Tilt series were acquired using a dose-symmetric tilt scheme ^62^ with a 3° step over a total range of −54 to +63°, relative to the lamella pre-tilt, yielding 40 tilt images per tilt series. The nominal defocus was set between −2.5 and - 4 µm. Each tilt image was exposed for 1.17 seconds at an electron dose of ∼3.63 e⁻/Å², giving a total electron dose of ∼145 e⁻/Å² per tilt series. Movies were recorded in EER format ^25^. A total of 349 tilt series were collected from 25 lamellae in a 31-hour session.

### Tomogram reconstruction of *S. cerevisiae*

Tilt series were processed directly at the super-resolution pixel size using WarpTools (Warp v2.0.0dev32) ^1,63^, as described for *D. melanogaster* unless stated otherwise. EER movies were rendered at 1.195 Å/px using relion_convert_to_tiff with an EER upsampling factor of 2, grouping every 20 EER frames to yield 18 fractions per tilt image. Motion correction and CTF estimation were carried out on a 1×1×18 motion grid and a 2×2×1 CTF grid, fitting the CTF between 40 and 8 Å over a defocus range of 2.5–4 µm. Tilt series stacks were generated, and poor-quality tilt images were removed as described for *D. melanogaster*. Tilt series were aligned with AreTomo2 (v1.0.0) ^64^ at 14.34 Å/px using an -AlignZ value of 3500 Å for all tilt series. The defocus hand was checked and flipped. 3D-CTF corrected tomograms were reconstructed at 14.34 Å/px. A total of 319 tomograms were of a sufficient quality to interpret and analyse.

### STA of *S. cerevisiae* ribosomes at the super-resolution pixel size

Particles were extracted as 3D subtomograms by WarpTools and STA was carried out in RELION (v4) ^68^, unless stated otherwise. Ribosomes were picked by template matching using PyTOM ^47^ on 3D-CTF corrected tomograms at 14.34 Å/px. To generate an initial reference, 2,091 ribosomes were manually picked in 20 tomograms. Particles were extracted at 14.34 Å/px and an initial model was generated *de novo* without imposed symmetry. Matching used the initial model as a template at 14.34 Å/px, using a template box of 32 voxels, a spherical mask and a particle diameter of 360 Å. Up to 6,000 candidate peaks were extracted per tomogram.

Particles were extracted at 14.34 Å/px in a box size of 32 voxels and refined in RELION until the Nyquist limit was reached. This process was repeated at 9.56 Å/px in a box of 48 voxels, and at 4.78 Å/px in a box size of 96 voxels. An M species was created from the 4.78 Å/px refinement, with the particles resampled to 1.195 Å/px, an initial low-pass filter of 10 Å, and C1 symmetry. Particle poses (refine_particles) and tilt series alignment (refine_imagewarp 1×1) were refined first. Per-tilt defocus (ctf_defocus, initially with ctf_defocusexhaustive) was added once the resolution reached 4.4 Å. A final round additionally refined volume warp (refine_volumewarp 4×4×2×10). The final ribosome map reached 3.18 Å from 190,883 particles in 310 tomograms, beyond the physical Nyquist limit of 4.78 Å. Resolution was determined by relion_postprocess using the M half-maps and a mask covering the whole ribosome (extended by 1 pixel with a soft edge of 8 pixels), with an FSC threshold of 0.143.

### Rosenthal–Henderson plots

After M refinement was completed, particle subsets were randomly selected and each subset was used to create a separate M species. Particles were sampled separately from each half-set to preserve the half-set assignments of the full dataset. Half-maps of each subset and of the full dataset were reconstructed in M without further refinement (--iter 0), retaining the particle poses and tomogram deformation parameters from the final M refinement of the maps with all particles present. Subset resolutions therefore reflect particle number at the alignment accuracy of the full dataset and exclude the effect of re-refining alignments from fewer particles. Resolutions were determined using the M half-maps with an FSC threshold of 0.143. To generate the Rosenthal-Henderson plot, 1/d² was plotted against the natural logarithm of the number of particles ^41^.

## Supplementary figures

**Supplementary Figure 1.**
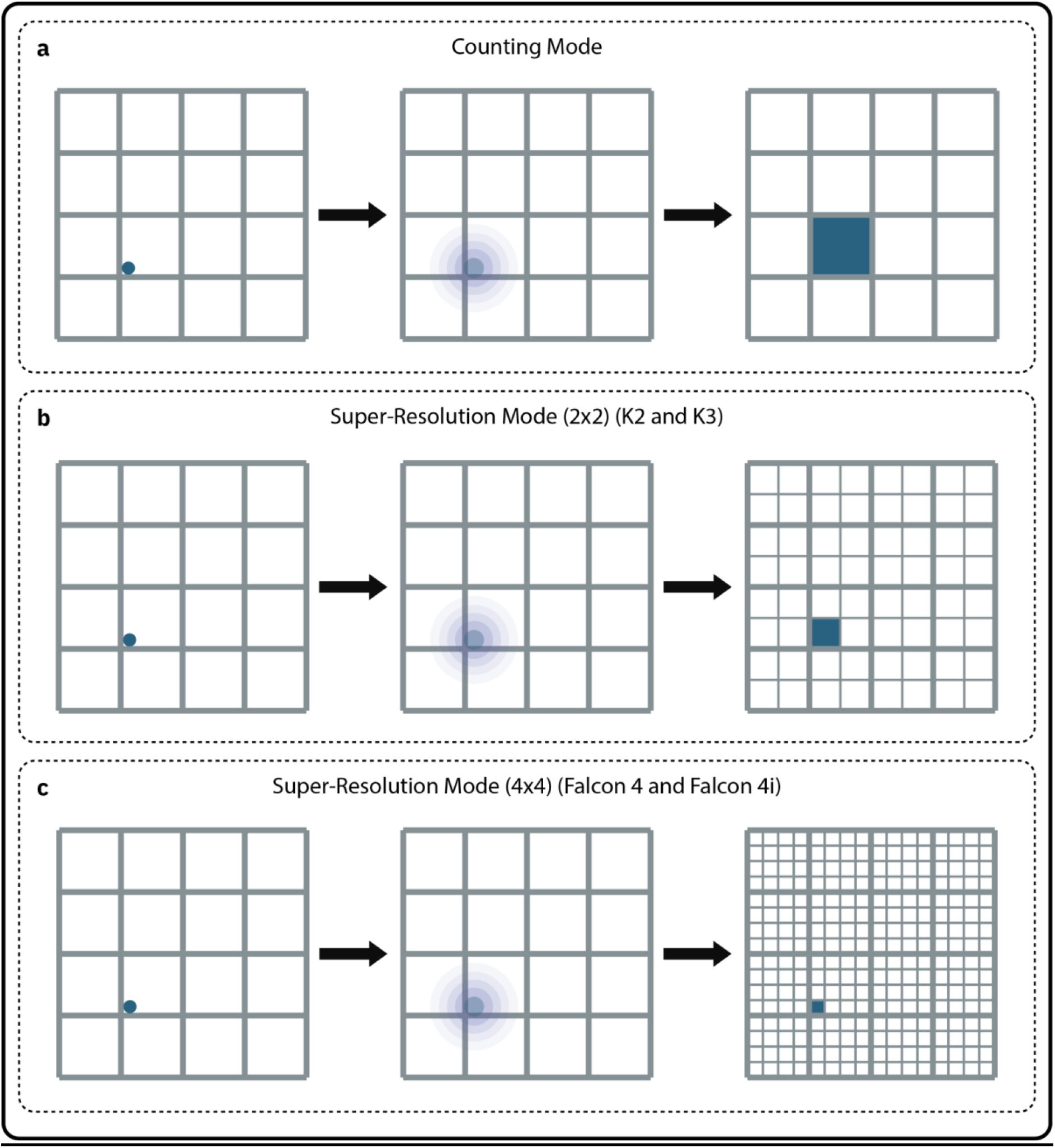
Assignment of electron events in counting and super-resolution processing. In all modes of processing, an electron hits a pixel on the detector (blue dot, left), the charge from this electron event diffuses across multiple pixels (blue cloud, middle), and the detector assigns signal to a particular pixel (blue pixel, right). (a) Counting mode: the point spread function of a single electron event spreads across neighbouring pixels, and a thresholding algorithm assigns the entire event to the pixel receiving the largest share. (b) Super-resolution mode on the K2 and K3 detectors: each physical pixel is subdivided into a 2 × 2 grid and a centroiding algorithm assigns the event to one quadrant. (c) The EER format on the Falcon 4 and 4i: each event is recorded on a 4 × 4 subpixel grid, and images are reconstructed on that grid and Fourier-cropped to the chosen pixel size at rendering. Figure partially generated with ChatGPT and is based on figure from Burton Smith *et al*. ^28^.

**Supplementary Figure 2.**
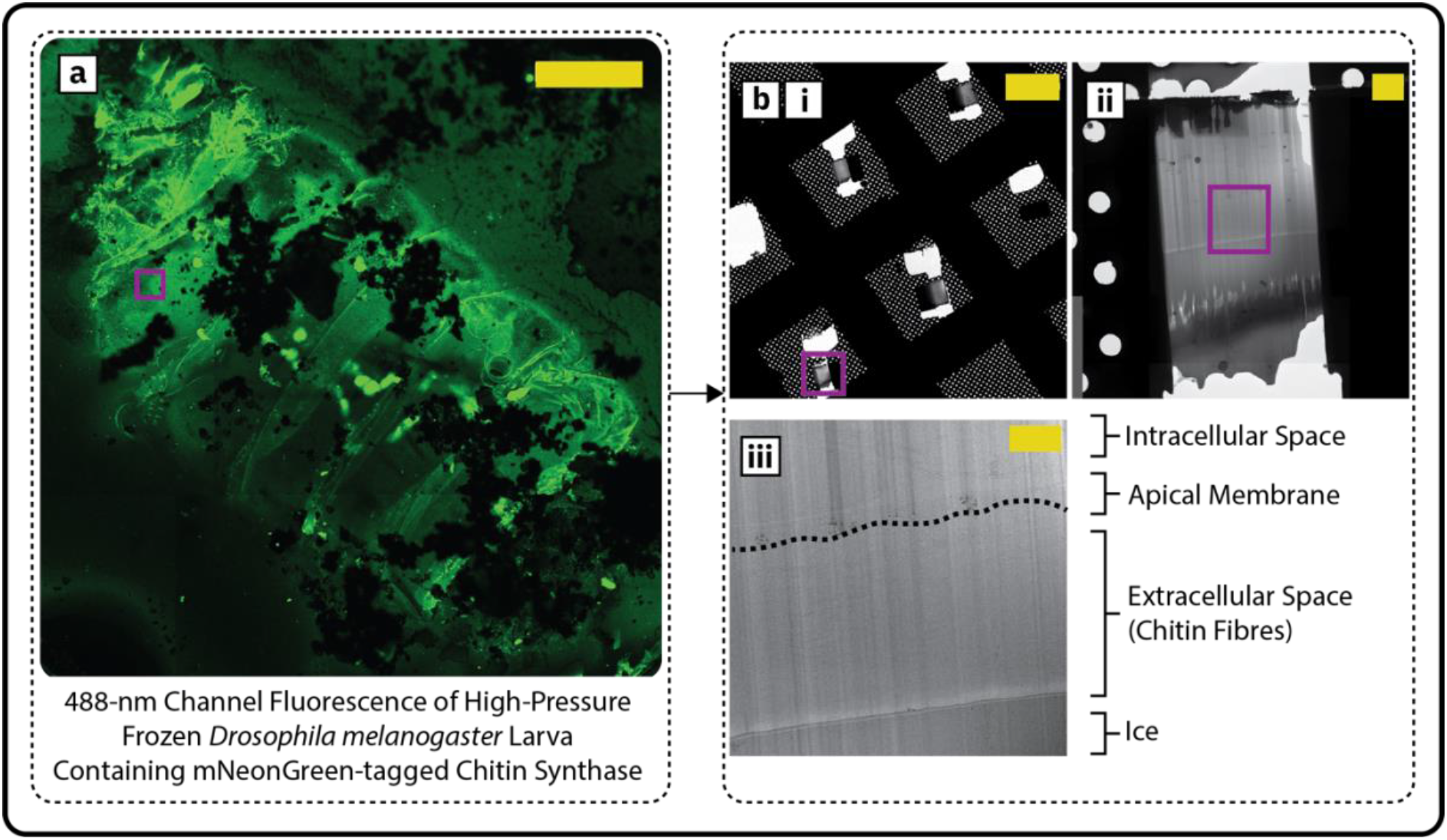
Targeting and serial lift-out of *Drosophila melanogaster* larva. (a) High-pressure frozen larva imaged by confocal microscopy. A 488-nm laser was used to acquire images of mNeonGreen-tagged chitin synthase signal. Most of the signal is due to specimen auto-fluorescence, which was used to morphologically map the larva. Chitin synthase localises to plaques on apical membrane undulae at the interface of the apical membrane and the cuticle ^31,32^. A region containing numerous fluorescent puncta was therefore targeted for serial lift-out and is highlighted in the purple box. Scale bar = 250 µm. (b) Images of lamellae taken in the transmission electron microscope. (i) Overview of the four lamellae generated from the lifted-out block. Scale bar = 50 µm. (ii) Overview of whole lamella highlighted in the purple box in *b(i)*. Scale bar = 2.5 µm. (iii) Close-up of the region highlighted in the purple box in *b(ii)*. The interface between the apical membrane and the cuticle (extracellular space) is traced with the dotted line. Tilt series acquisition was mostly focused on this interface. Scale bar = 1 µm.

**Supplementary Figure 3.**
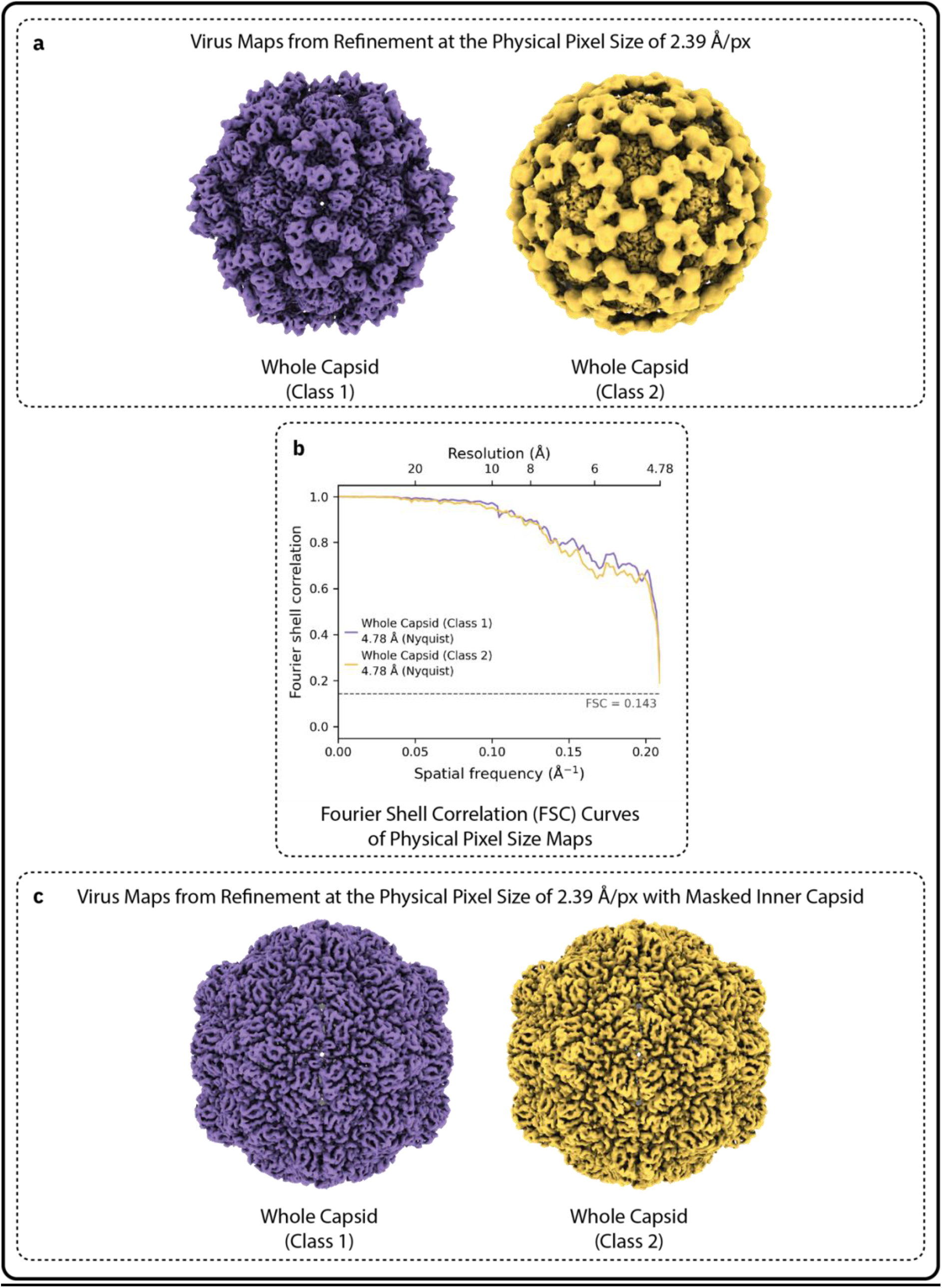
Comparison of Class 1 and Class 2 at physical pixel size. (a) Virus maps processed at physical pixel size (2.39 Å/px). Class 1 is coloured purple and Class 2 is coloured yellow. Both maps are shown in the same orientation. (b) FSC curves for the reconstructions in *a*, with the 0.143 threshold marked. (c) Comparison of the inner capsids of Class 1 and Class 2. Both maps are masked by the same inner capsid mask and are shown at the same orientation and threshold. Maps shown in *a* and *c* were denoised by M.

**Supplementary Figure 4.**
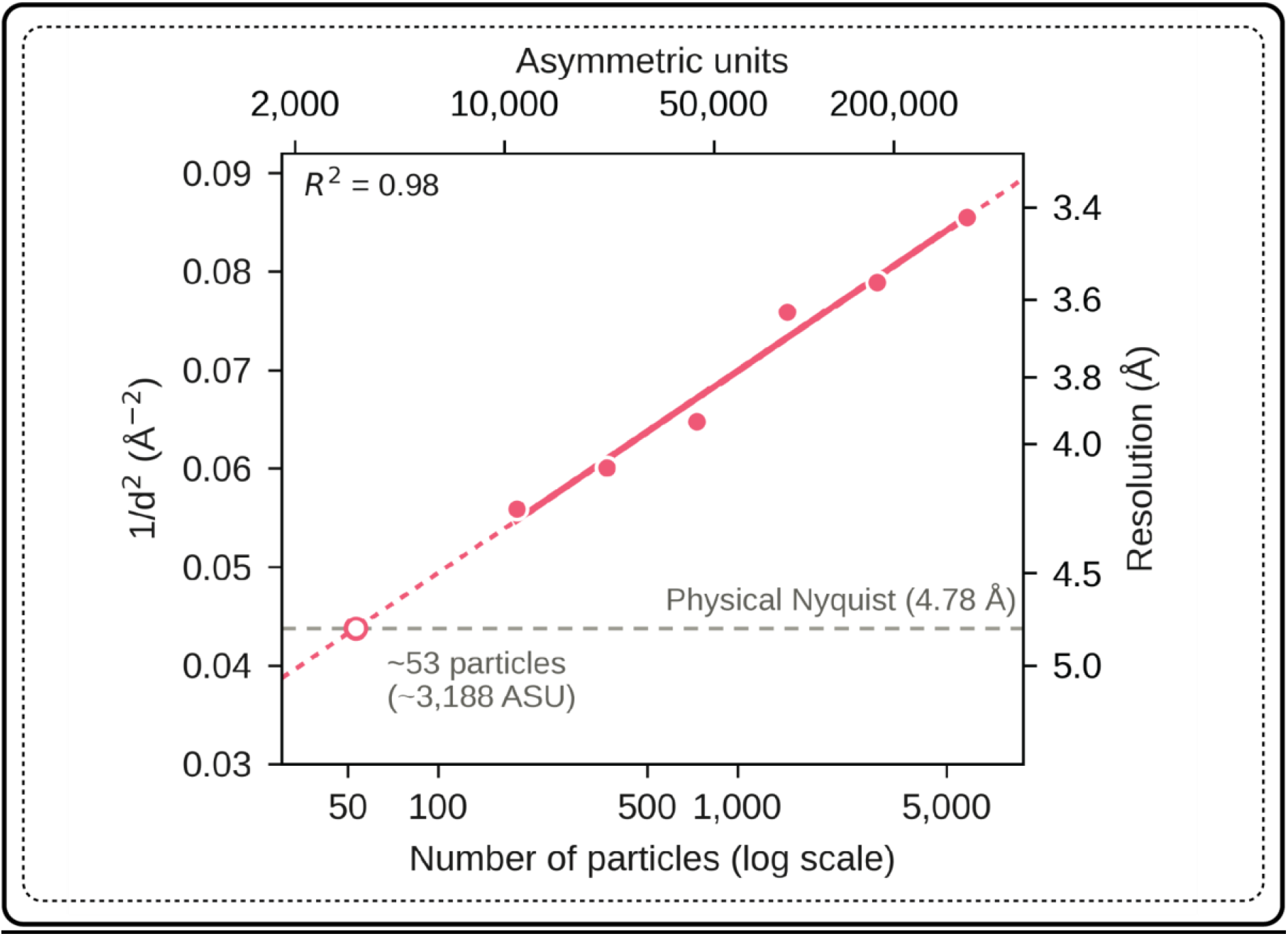
Rosenthal–Henderson plot of the combined inner capsid reconstruction. B-factor plot showing the number of particles required to exceed the physical Nyquist limit of 4.78 Å.

**Supplementary Figure 5.**
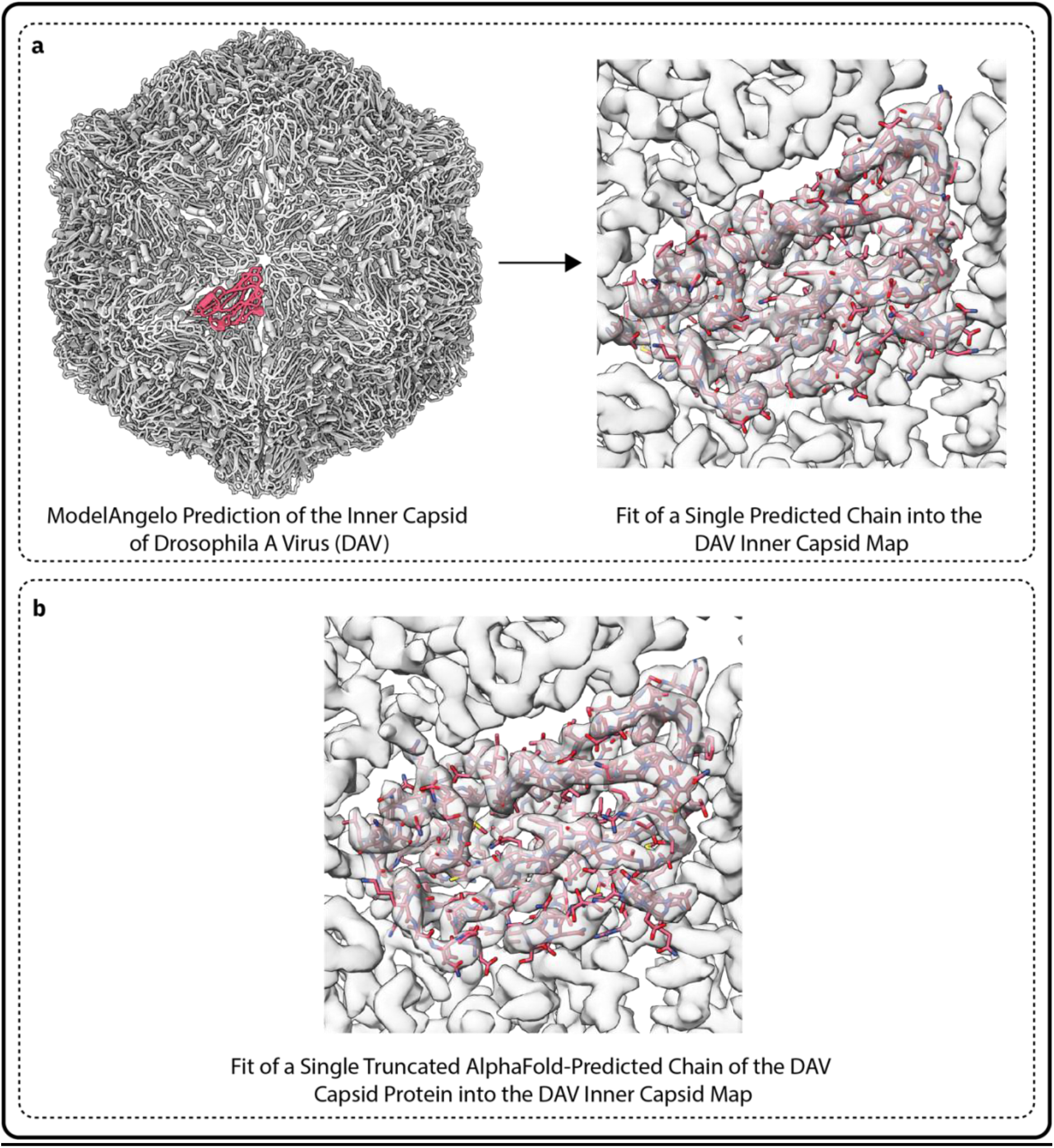
*De novo* sequencing using ModelAngelo and AlphaFold. (a) An unrefined model of the DAV inner capsid, as predicted by ModelAngelo. No prior sequence was given to ModelAngelo. All chains are coloured grey, except for one 169 residue chain which is coloured red. The sequence of this chain was submitted to BLAST. The fit of the chain into the inner capsid map is displayed on the right. (b) The AlphaFold prediction of the structure of the DAV capsid protein, truncated to only include inner capsid residues. The structure was rigid-body fitted into the same region of the map as shown in *a* (main chain map-model CC = 0.69). Maps shown in *a* and *b* were denoised by M.

## Supplementary videos

**Supplementary Video 1. Virus-containing tomograms.** Tomographic slices through several tomograms showing the range of particle arrangements, from isolated particles to tightly packed lattice arrays. Scale bar = 200 nm.

**Supplementary Video 2. Class distribution of viruses in the tomograms.** Class 1 (purple) and Class 2 (yellow) virus particle positions back-plotted into several tomograms, highlighting their distribution between the intracellular space and the cuticle. Note that particles that appear unpicked were classified as junk or lower quality particles and removed from the dataset. Scale bar = 200 nm.

## Supplementary tables

**Supplementary Table 1:** Conditions for cryo-electron tomography data collection.

|  | *Drosophila melanogaster* (virus) | *Saccharomyces cerevisiae* (ribosome) |
| --- | --- | --- |
| Data collection |  |  |
| Microscope | Titan Krios G4 | Titan Krios G4 |
| Detector | Falcon 4i | Falcon 4i |
| Voltage (kV) | 300 | 300 |
| Spherical aberration (mm) | 2.7 | 2.7 |
| Energy filter slit (eV) | 10 | 10 |
| C2 aperture (μm) | 50 | 50 |
| Objective aperture (μm) | 70 | 70 |
| Acquisition software | SerialEM v4.2.5 / PACE-tomo | SerialEM v4.3.0beta / PACE-tomo |
| Nominal magnification | 53,000× | 53,000× |
| Calibrated physical pixel size (Å) | 2.39 | 2.39 |
| Super-resolution pixel size (Å) | 1.195 | 1.195 |
| Movie format | EER | EER |
| EER hardware frames per tilt movie | 351 | 360 |
| EER frame grouping (fractions per tilt image) | 25 (14 fractions) | 20 (18 fractions) |
| Nominal defocus range (μm) | −3 to −6 | −2.5 to −4 |
| Tilt scheme | Dose-symmetric | Dose-symmetric |
| Tilt range relative to lamella pre-tilt (°) | −51 to +51 | −54 to +63 |
| Tilt step (°) | 3 | 3 |
| Fluence per tilt image (e<sup>−</sup>/Å<sup>2</sup>) | ~3.65 | ~3.63 |
| Total fluence (e<sup>−</sup>/Å<sup>2</sup>) | ~128 | ~145 |
| Lamellae | 4 | 25 |
| Tilt series collected | 32 | 349 |
| Tomograms reconstructed | 28 | 319 |
| Tomograms contributing particles | 20 | 310 |

**Supplementary Table 2:** Processing parameters and deposition information for the subtomogram averages.

|  | DAV Class 1 | DAV Class 2 | DAV inner capsid | S. cerevisiae ribosome |
| --- | --- | --- | --- | --- |
| **Processing** |  |  |  |  |
| **Tilt series alignment** | AreTomo2 v1.0.0 | AreTomo2 v1.0.0 | AreTomo2 v1.0.0 | AreTomo2 v1.0.0 |
| **Tomogram reconstruction** | WarpTools (Warp v2.0.0dev36) | WarpTools (Warp v2.0.0dev36) | WarpTools (Warp v2.0.0dev36) | WarpTools (Warp v2.0.0dev32) |
| **Subtomogram averaging software** | RELION v4 / M (Warp v2.0.0dev36) | RELION v4 / M (Warp v2.0.0dev36) | RELION v4 / M (Warp v2.0.0dev36) | RELION v4 / M (Warp v2.0.0dev32) |
| **Symmetry imposed on final map** | I2 | I2 | I2 | C1 |
| **Particles** | 3,513 | 2,342 | 5,855 | 190,883 |
| **Asymmetric units (60 per particle)** | 210,780 | 140,520 | 351,300 | n/a |
| **Mask used for FSC** | Whole capsid | Whole capsid | Inner capsid | Whole ribosome |
| **Resolution (Å) (FSC = 0.143)** | 3.94 | 4.08 | 3.42 | 3.18 |
| **Map sharpening B-factor (Å<sup>2</sup>)** | -64.6 | -66.5 | -53.7 | -29.2 |
| **EMDB accession** | EMD-XXX | EMD-XXX | EMD-XXX | EMD-XXX |

**Supplementary Table 3:** Model building, refinement, and composition of the Drosophila A virus inner capsid model. Statistics are reported for the asymmetric unit.

|  | DAV inner capsid |
| --- | --- |
| **Model building** |  |
| **Map used for model building** | Super-resolution processed map of the DAV inner capsid, 3.42 Å (sharpened) |
| **Sequence-primed initial model building** | ModelAngelo v1.0.9 with DAV capsid protein sequence |
| **Manual rebuilding** | Coot v0.9.7 |
| **Refinement** |  |
| **Refinement package** | Phenix v2.1 (phenix.real_space_refine) |
| **Resolution used for refinement (Å)** | 3.5 |
| **Refinement rounds (Phenix / Coot cycles)** | 4 |
| **Symmetry** | Icosahedral (I2); neighbours regenerated with phenix.apply_ncs |
| **Model composition** |  |
| **Chains (asymmetric unit)** | 3 (BB, BC, BD) |
| **Residue range modelled** | 69–240 (172 residues per chain) |
| **Non-hydrogen atoms** | 3,903 |

**Supplementary Table 4:**
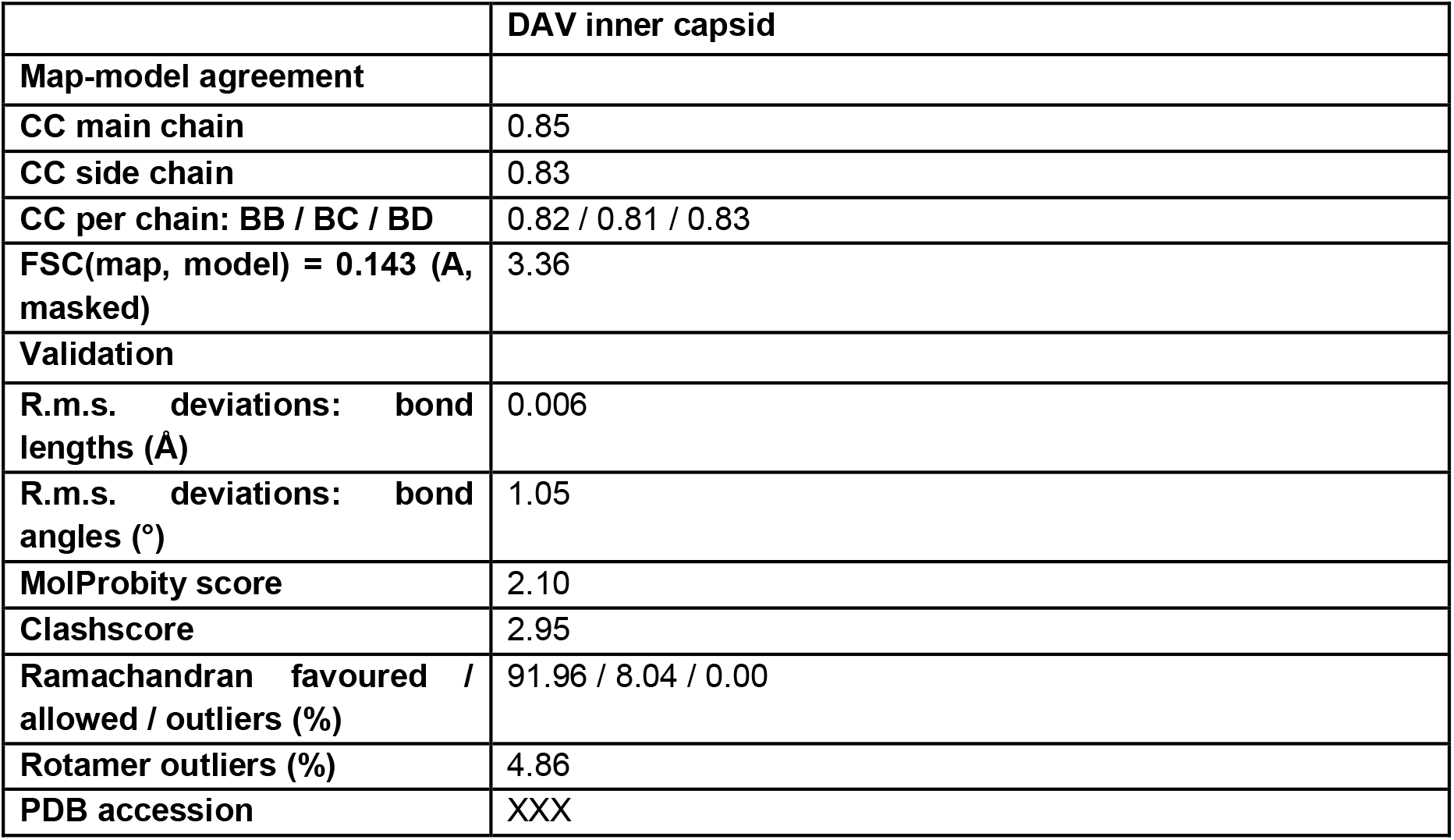
Validation statistics and deposition information for the Drosophila A virus inner capsid model. Statistics are reported for the asymmetric unit.

## Supplementary scripts

**Supplementary Script 1.** Script used to calculate the real-space correlation coefficient between the Class 1 and Class 2 half-maps at the physical pixel size. Usage: python3 map_cc.py map1.mrc map2.mrc mask.mrc [threshold_fraction].

