## supplementary_scripts for "Super-Resolution Processing Enables *In Situ* Subtomogram Averaging Beyond the Nyquist Limit": map_cc.py

```
#!/usr/bin/env python3
"""Real-space correlation between two MRC maps inside a mask.

Usage:  python3 map_cc.py map1.mrc map2.mrc mask.mrc [threshold_fraction]

threshold_fraction (default 0.5) is applied relative to the mask's maximum
value, so soft masks that never reach 1.0 are handled correctly.
Reads the maps in slabs, so memory use stays low.
"""
import sys, numpy as np

def header(path):
    h = np.fromfile(path, dtype=np.int32, count=256)
    hf = h.view(np.float32)
    nx, ny, nz, mode = int(h[0]), int(h[1]), int(h[2]), int(h[3])
    if mode != 2:
        sys.exit('%s: only mode 2 (float32) supported, got mode %d' % (path, mode))
    return nx, ny, nz, 1024 + int(h[23])

def slab(path, off, nx, ny, z0, z1):
    return np.fromfile(path, dtype=np.float32, count=(z1 - z0) * ny * nx,
                       offset=off + z0 * ny * nx * 4).astype(np.float64)

def main(f1, f2, fm, frac=0.5, chunk=32):
    g = [header(f) for f in (f1, f2, fm)]
    if len({x[:3] for x in g}) != 1:
        sys.exit('maps have different dimensions: %s' % [x[:3] for x in g])
    nx, ny, nz = g[0][:3]
    # mask maximum, measured rather than taken from the header (headers go stale)
    mmax = max(float(slab(fm, g[2][3], nx, ny, z0, min(z0 + chunk, nz)).max())
               for z0 in range(0, nz, chunk))
    if not mmax > 0:
        sys.exit('mask maximum is %g' % mmax)
    thr = frac * mmax
    S = dict.fromkeys(('n','sx','sy','sxx','syy','sxy','wn','wsx','wsy','wsxx','wsyy','wsxy'), 0.0)
    for z0 in range(0, nz, chunk):
        z1 = min(z0 + chunk, nz)
        a = slab(f1, g[0][3], nx, ny, z0, z1)
        b = slab(f2, g[1][3], nx, ny, z0, z1)
        m = slab(fm, g[2][3], nx, ny, z0, z1)
        sel = m > thr
        x, y = a[sel], b[sel]
        w = m / mmax
        S['n'] += x.size;       S['sx'] += x.sum();         S['sy'] += y.sum()
        S['sxx'] += (x*x).sum(); S['syy'] += (y*y).sum();   S['sxy'] += (x*y).sum()
        S['wn'] += w.sum();     S['wsx'] += (w*a).sum();    S['wsy'] += (w*b).sum()
        S['wsxx'] += (w*a*a).sum(); S['wsyy'] += (w*b*b).sum(); S['wsxy'] += (w*a*b).sum()
    def cc(n, sx, sy, sxx, syy, sxy):
        return (sxy - sx*sy/n) / np.sqrt((sxx - sx*sx/n) * (syy - sy*sy/n))
    if S['n'] == 0:
        sys.exit('no voxels above threshold %.4g (mask max %.4g)' % (thr, mmax))
    print('mask maximum          : %.4g' % mmax)
    print('threshold             : %.4g  (%.2f x max)' % (thr, frac))
    print('voxels inside mask    : %d  (%.2f %% of box)' % (S['n'], 100*S['n']/(nx*ny*nz)))
    print('CC inside mask        : %.4f' % cc(S['n'], S['sx'], S['sy'], S['sxx'], S['syy'], S['sxy']))
    print('CC mask-weighted      : %.4f' % cc(S['wn'], S['wsx'], S['wsy'], S['wsxx'], S['wsyy'], S['wsxy']))

if __name__ == '__main__':
    if len(sys.argv) < 4:
        sys.exit(__doc__)
    main(sys.argv[1], sys.argv[2], sys.argv[3],
         float(sys.argv[4]) if len(sys.argv) > 4 else 0.5)
```
